# Multi-trait evaluation of a tomato MAGIC population identifies promising lines with improved nitrogen use efficiency (NUE)

**DOI:** 10.64898/2026.07.14.738388

**Authors:** Virginia Baraja-Fonseca, Daniel Gil-Villar, Jon Bančič, Begoña Renau-Morata, María Salud Justamante, Mariola Plazas, Pietro Gramazio, Santiago Vilanova, José Manuel Pérez-Pérez, Antonio Granell, Rosa Victoria Molina, Sergio G. Nebauer, Jaime Prohens, Andrea Arrones

**Affiliations:** Instituto de Conservación y Mejora de la Agrodiversidad Valenciana, Universitat Politècnica de València, Camino de Vera s/n, 46022, València, Spain; Departamento de Producción Vegetal, Universitat Politècnica de València, Camino de Vera s/n, 46022, València, Spain; Instituto Universitario de Biotecnología y Biomedicina, Departamento de Biología Vegetal, Universitat de València, Vicent Andrés Estellés 19, 46100, Burjassot, València, Spain; Instituto de Bioingeniería, Universidad Miguel Hernández, Avda. de la Universidad s/n, 03202, Elche, Spain; Instituto de Biología Molecular y Celular de Plantas, Consejo Superior de Investigaciones Científicas, Universitat Politècnica de València (IBMCP-CSIC-UPV), València, 46022, Spain

**Keywords:** tomato breeding, MAGIC, nitrogen use efficiency, transgressive segregation, selection indices, line selection

## Abstract

Nitrogen-use efficiency (NUE) is a pivotal breeding target in tomato (*Solanum lycopersicum* L.) to sustain production under reduced N inputs. Here, we leveraged a recently developed tomato multi-parent advanced generation inter-cross (ToMAGIC) population to identify lines with superior performance under reduced N availability. The eight founders and a core subset of 118 ToMAGIC lines were characterized with 10,684 SNP markers and evaluated under optimal (opN, 15 mM) and suboptimal (subN, 8 mM) N supply in an experiment totalling 1,576 plants, generating 48,068 data points across 61 phenotypic variables. Under both N treatments, ToMAGIC lines exhibited transgressive segregation for most traits, confirming the value of this population as a reservoir of untapped variation. Notably, under subN conditions, harvest index (Hi) increased by 29-44%, suggesting adaptive resource redistribution toward reproductive sinks. Variance partitioning revealed that agronomic and NUE-related traits were largely under genetic control, with heritability estimates frequently above 0.80 and broadly conserved across N treatments. Multivariate trait analysis identified fruit yield N concentration (NUE component, C_N,y_), shoot biomass N content (NAb), and shoot growth-related traits as the main drivers of treatment differentiation. Finally, proxy traits were prioritized by integrating response magnitude, heritability, trait correlations, and treatment-discriminatory power into multi-trait selection indices. This strategy generated favorable predicted genetic gains, reaching 158% for high-performance lines and 170% for subN-adapted lines, and consistently identified lines 402, 428, 518, 800, and 816 as promising pre-breeding materials. Overall, this study supports ToMAGIC as a powerful resource for developing N-efficient cultivars suited for sustainable agriculture.

## Introduction

Tomato (*Solanum lycopersicum* L.) is the most economically important vegetable crop, with global production exceeding 185 million tons annually (FAOSTAT, 2025). Tomato cultivation, like other vegetable crops, relies heavily on fertilizer inputs, particularly nitrogen (N), to ensure high yields and fruit quality (Cheng et al., 2021; Adalibieke et al., 2023). However, a substantial proportion of applied N is not utilized by plants and is instead lost through processes such as leaching, volatilization, surface runoff, and denitrification (Wang and Li, 2019; Govindasamy et al., 2023). These losses contribute to environmental pollution and reduce fertilizer efficiency. In response, many international policies aim to decrease nutrient losses and promote more sustainable fertilization practices. For instance, within the Farm to Fork Strategy, the European Commission has set targets to reduce nutrient losses by at least 50% while ensuring no deterioration in soil fertility, and to reduce fertilizer use by at least 20% by 2030 (https://agriculture.ec.europa.eu). Reaching these objectives will require developing tomato varieties with improved N use efficiency (NUE), enabling reduced reliance on external N inputs during cultivation.

From a breeding perspective, improving NUE involves developing genotypes capable of capturing and utilizing N more efficiently under N-limited conditions while maintaining high yield. A great effort has been made in field crops such as barley, maize, rice, and wheat, where cultivar differences and breeding gains under contrasting N supply have been documented (Laidig et al., 2024; Natarajan et al., 2025; Huang et al., 2025; Saenz et al., 2025). In vegetable crops, specifically in tomato, however, NUE-oriented breeding remains comparatively underdeveloped. This may be partly due to the relatively narrow genetic base of elite germplasm, shaped by domestication and breeding bottlenecks (Blanca et al., 2015). Nevertheless, recent studies have revealed substantial variability among tomato landraces, old cultivars, and wild relatives in their response to low N availability (Tripodi et al., 2022; Flores-Saavedra et al., 2024; Machado et al., 2024; Renau-Morata et al., 2024; Cirillo et al., 2025), highlighting the potential of broadening the germplasm base to develop more N-efficient cultivars.

To capture and exploit this diversity, our research group developed a multi-parent advanced generation inter-cross (MAGIC) population using eight genetically diverse founder accessions, including four *S. lycopersicum* var. *cerasiforme* (SLC) and four *S. pimpinellifolium* (SP) accessions (ToMAGIC, Arrones et al., 2024). The potential of this population is reinforced by the prior characterization of its founders, whose whole-genome resequencing and morphoagronomic evaluation revealed extensive diversity and a large number of private variants, highlighting their value as a source of novel alleles for broadening the genetic base of tomato breeding (Gramazio et al., 2020). The resulting ToMAGIC population, comprising 345 inbred lines, encompasses broad genetic diversity resulting from admixture and recombination of founder alleles and shows extensive phenotypic variation in agronomic traits (Arrones et al., 2024). In addition, recent evaluation of the SLC and SP founders revealed marked variation in N partitioning and overall NUE under reduced N inputs (Gil-Villar et al., 2025). Taken together, these findings highlight the potential of this genetic background for breeding tomato genotypes better adapted to suboptimal N supply and provide a strong rationale for evaluating the ToMAGIC population to identify lines combining improved NUE with favorable agronomic performance.

In this study, the eight founder accessions and a core subset of 118 ToMAGIC lines were evaluated under optimal (opN, 15 mM) and suboptimal (subN, 8 mM) N supply conditions. This core subset was designed to represent the genetic diversity of the full ToMAGIC population based on molecular marker data (Arrones et al., 2024). The subN treatment represents a substantial N reduction, potentially contributing to policy goals aimed at reducing nutrient losses and fertilizer dependence. These materials were characterized for a broad set of above- and below-ground agronomic and physiological traits associated with NUE to identify informative phenotypic indicators for NUE selection. Phenotypic and genomic information was then integrated through mixed-model analyses to estimate genomic breeding values (GEBVs), which were used in multi-trait selection indices to rank genotypes according to two breeding objectives: (i) the identification of high-performance lines with strong overall agronomic performance and (ii) the identification of lines specifically adapted to subN conditions. This integrative approach enabled the identification of tomato lines with improved performance under reduced N inputs and provided insights into the phenotypic traits most informative for NUE-oriented breeding.

## Results

### Phenotypic variation across ToMAGIC lines under opN

To assess phenotypic variation relevant to NUE, the core subset of 118 ToMAGIC lines, together with the eight founder accessions, was evaluated under greenhouse conditions. Overall, the experiment comprised 1,576 plants and generated a large dataset from repeated physiological measurements, destructive biomass samplings, fruit production records, root architecture traits, and C and N elemental analyses, resulting in 61 phenotypic variables and 48,068 data points (Tables S1 and S2). Under opN conditions, substantial phenotypic variation was observed across the ToMAGIC core subset (Table S3), indicating broad diversity for traits potentially contributing to NUE. Specifically, yield-related traits showed the highest coefficients of variation (CV > 0.5), including fruit production (Prod) and harvest index (Hi), as well as for NUE-related traits such as fruit C (CFr) and N (NFr) contents, yield-specific N efficiency (E_N,y_) and NUE itself (Table S3). By contrast, physiological parameters like maximum efficiency of PSII (Fv/Fm), normalized difference vegetation index (NDVI), leaf anthocyanins (Ant) and flavonols (Flv) contents, and the average root diameter (Rdi) were among the least variable traits, with CV values below 0.1 (Table S3).

Several lines exhibited vigorous plant architecture, with the ToMAGIC lines showing an average shoot biomass (Ab) of 675 g fresh weight (FW) at the end of the experiment (135 days-old; T5), with no significant differences from either founder group (Table S3). This vigorous architecture was complemented by favorable reproductive performance. No significant differences among groups were detected for Prod, fruit count (Frc), or Hi, with the ToMAGIC subset showing mean values of 107 g FW and 10 g dry weight (DW) for Prod, 53 fruits per plant for Frc, and 0.12 FW and 0.08 DW for Hi. By contrast, significant differences were observed for average fruit weight (Frw). In FW, ToMAGIC lines showed an intermediate mean Frw between SLC and SP founders (2.09, 2.96, and 1.11 g, respectively), whereas in DW they were closer to the SLC accessions (0.20 vs 0.21 g) (Table S3). Physiological traits did not show significant differences among groups. Regarding NUE-related traits, mean NUE and N uptake efficiency (U_N_) values in the ToMAGIC subset (1,752 and 375 g/g, respectively) were comparable to those of the SP founders. No significant differences were detected for C and N contents. In the ToMAGIC lines, shoot biomass contained on average approximately 14 times more C (CAb) than N (NAb) (49.04 vs 3.46 g/plant), whereas fruits contained approximately 17 times more C than N (5.18 vs 0.29 g/plant) (Table S3).

In particular, the presence of lines with high Prod, Hi, CFr, NFr, and E_N,y_ indicates that part of the population showed efficient C and N partitioning towards fruits (Table S3). Thus, beyond total biomass accumulation, the ToMAGIC collection showed relevant variation in partitioning for yield, a key component for selecting lines that maintain productivity and NUE under reduced N supply.

### Phenotypic responses to N supply limitation

Assessing the performance of the core subset of ToMAGIC lines under N-limited conditions is crucial for NUE improvement. To this end, lines were also assayed under suboptimal (subN, 8 mM N) N supply. Compared with performance under opN treatment (15 mM N), significant phenotypic deviations were observed in the ToMAGIC lines, highlighting the impact of N limitation on key traits (Figure 1A). In particular, biomass-related traits, including total plant biomass (Tb), vegetative plant biomass (Plb), root biomass (Rb), and Ab, were significantly reduced under subN at T5, declining by approximately one-third in both FW and DW, except for Rb, which showed a smaller reduction of nearly 20% (Figure 1A and Table S3). Regarding yield-related traits, Prod remained relatively stable, with only moderate decreases (8.70% FW and 0.93% DW), whereas Hi increased by 29% FW and 44% DW, indicating a shift towards greater biomass partitioning to reproductive sinks under reduced N supply (Figure 1A). However, despite these overall shifts, substantial phenotypic variation among lines was maintained for most traits (Table S3). Lines remained vigorous and productive under subN, with average values of 439 g FW for Ab at T5, 48 fruits per plant, and 98 g FW for Prod. Transgressive segregation was also observed for these traits, with 3-8% of lines showing positive transgression, while negative transgression was more frequent for Ab_FW_T5 and Frc, affecting 17% and 14% of lines, respectively (Table S4). Consistently, the highest values under subN were observed in line 508 for Ab at T5 (660 g FW), line 800 for Frc (122 fruits per plant), and line 428 for Prod (410 g FW) (Tables S2 and S4).

**Figure 1.**
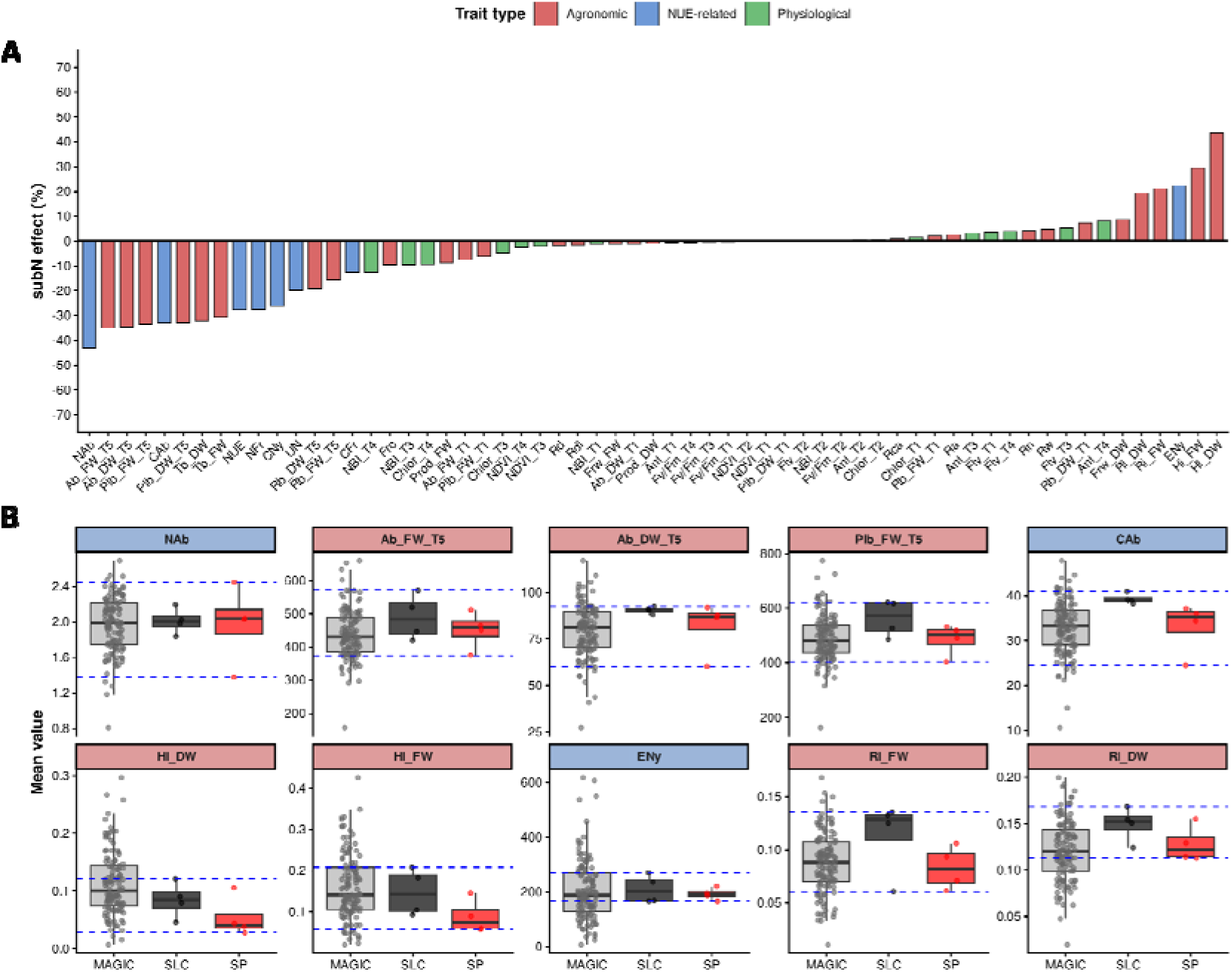
Phenotypic variation across the core subset of ToMAGIC lines. **A.** Relative effect of suboptimal nitrogen supply (subN; 8 mM N) on all the evaluated traits. Bars represent the percentage change in trait mean values under subN relative to optimal nitrogen supply (opN; 15 mM N). Traits were defined in Table S1. **B.** Distribution of the five most positively and five most negatively affected traits under subN supply in the ToMAGIC core subset relative to the *S. lycopersicum* var. *cerasiforme* (SLC) and *S. pimpinellifolium* (SP) founders. Points correspond to individual genotype means. Blue dashed lines indicate the phenotypic range defined by the founder accessions. No significant differences among groups were detected for these traits based on Tukey-adjusted pairwise comparisons of estimated marginal means from mixed models.

Furthermore, the root index (Ri) increased by 21% FW and 19% DW, likely supported by a slight increase in most root-related traits under limited N availability. Over time, the leaf chlorophyll content (Chlor), the nitrogen balance index (NBI), and NDVI were slightly negatively affected under subN, whereas Flv and Ant showed slight increases (Figure 1B). For NUE-related traits, reduced N availability decreased NUE by 28%, mainly through changes in its U_N_ and C_N,y_ components. However, NUE still displayed a range wider than that observed in the founders, with line 856 showing the highest value (3,653 g/g) and line 418 the lowest value (39 g/g) (Table S3). Nevertheless, E_N,y_ improved by 22%, suggesting a more efficient N redistribution toward improving yield (Figure 1B).

Interestingly, several transgressive lines showed favorable combinations of traits under subN, particularly for reproductive performance and NUE-related parameters (Table S4). Line 428 was especially notable, achieving the highest Prod value and showing positive transgressions for Frc, Hi, E_N,y_, NUE, NFr, and CFr, along with several root architecture traits. Line 800 also showed a strong reproductive profile, with the highest Frc and positive transgression for Prod, Hi, E_N,y_, NFr, and CFr (Table S4). Other lines, such as 518 and 816, displayed positive transgression for traits related to fruit production, biomass partitioning, NUE, and C and N allocation to fruits. In contrast, lines 508 and 761 were mainly transgressive for biomass-related traits, suggesting distinct phenotypic strategies within the population (Table S4). Together, these results indicate that the ToMAGIC population contains lines with distinct but favorable responses to reduced N supply, including genotypes with enhanced partitioning for yield and NUE-related performance.

### Trait analysis under opN and subN: effect, variance, and heritability

To determine whether the phenotypic differences observed between N treatments within the core subset of ToMAGIC lines were associated with genetic variation and treatment-specific responses, each trait was analyzed using a multi-treatment mixed model (see Materials and Methods). Significant effects were detected for all sources of variation included in the model (Table S5). The N treatment effect was generally not significant at early stages (T1-T2), except for Rb and Fv/Fm. However, the effect of N availability became evident at later time points, suggesting an intensified stress effect with prolonged exposure to the treatment (Table S5). For all traits except root depth (Rd), the genotype random effect structured by N explained a significant proportion of the variation, revealing significant genetic variability in trait expression in at least one N condition (Figure 2A and Table S5). On the other hand, spatial adjustments were required for all traits, and for some of them, model fit improved when heterogeneous residual variances were considered (Table S5).

**Figure 2.**
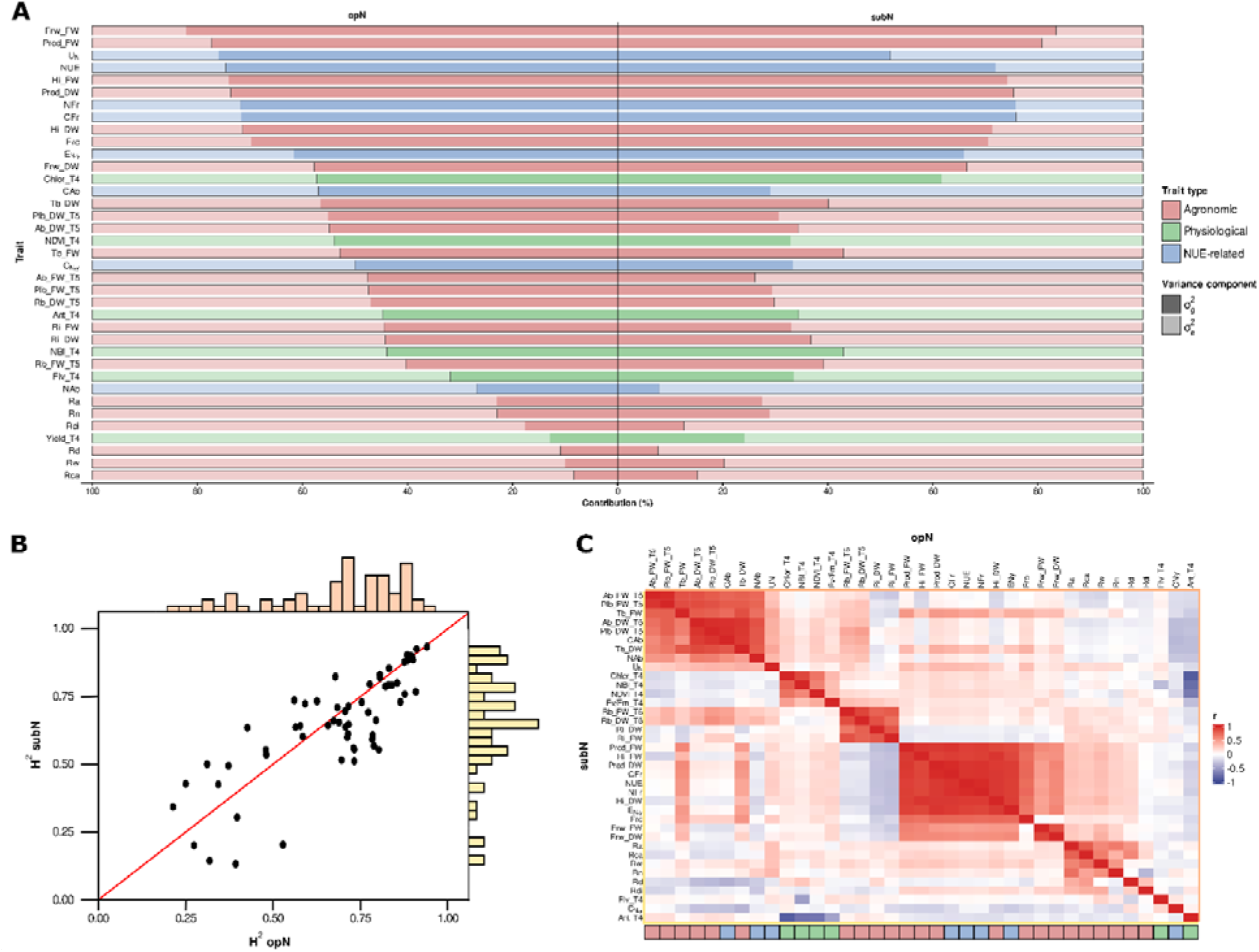
Genetic parameters and genotypic correlations estimated from the diagonal multi-treatment mixed model across the core subset of ToMAGIC lines under optimal nitrogen (opN; 15 mM N) and suboptimal nitrogen (subN; 8 mM N) supply. **A.** Relative contribution (%) of additive genetic (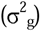) and residual (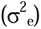) variance components for traits measured at final time points. **B.** Correlation between narrow-sense heritability (h^2^) estimates across N levels. The red line represents the 1:1 relationship, and marginal histograms show the distribution of heritability values for each N level. **C.** Empirical genotypic trait correlations based on best linear unbiased predictions (BLUPs).

Variance component and heritability estimates further clarified the sources of variation underlying trait expression under opN and subN conditions. Several traits, including some associated with shoot biomass, fruit production, physiology, and NUE, showed a predominance of genetic variance together with moderate-to-high heritability, indicating substantial genetic control over phenotypic variation (Figure 2A and Table S6). Under opN, the highest proportions of genetic variance were observed for average fruit weight (Frw_FW), Prod_FW and U_N_, accounting for 82%, 77%, and 76 % of the total variance, respectively (Figure 2A). These traits also showed high heritability estimates, with h^2^ = 0.94 for Frw_FW and h^2^ = 0.91 for both Prod_FW and U_N_ (Table S6). Under subN, both yield-related traits remained among those with the strongest genetic contribution, followed by CFr and NFr, which showed genetic variance proportions of 76% in both cases, and heritability estimates of h^2^ = 0.90 (Figure 2A and Table S6). By contrast, root-related and Fv/Fm traits showed lower genetic variance, larger residual components, and lower heritability estimates, with the lowest values observed for root convex area (Rca) under opN (h^2^ = 0.21) and Fv/Fm_T3 under subN (h^2^ = 0.14) (Figure 2A and Table S6). Heritability estimates were broadly comparable between N levels (r = 0.81), with mean values of 0.68 under opN and 0.65 under subN (Figure 2B).

### Relationships among traits under both N levels

Patterns of association among traits were investigated to characterize relationships among agronomic, physiological, and NUE-related variables within the core subset of ToMAGIC lines under opN and subN conditions. This analysis was performed using traits measured at the final time points, when the N treatment effect was most evident. Earlier time points were not included to avoid overrepresenting repeated measurements of the same trait, which were expected to provide largely overlapping information. This analysis helped reveal coordinated trait responses and distinguish complementary from redundant sources of information for selection. Several groups of traits displayed strong positive or negative genotypic correlations, forming clear clusters that were largely preserved across both N levels (r = 0.91) (Figure 2C). The overall correlation patterns were also highly consistent between phenotypic correlations estimated from raw data (Figure S1 and Table S7) and genotypic correlations derived from best linear unbiased predictions (BLUPs) (Figure 2C and Table S8), with correlations of 0.98 under both opN and subN.

Five main clusters of correlated traits could be distinguished: (i) plant biomass-related traits, (ii) root biomass-related traits, (iii) physiological traits, (iv) yield-related traits, and (v) root architecture-related traits (Figure 2C). The plant biomass cluster was strongly and positively correlated with NUE-related traits such as CAb and NAb contents, and U_N_, but negatively correlated with C_N,y_ and Ant. Root biomass traits, particularly Ri, showed weak negative correlations with yield-related traits. Among physiological traits, Chlor, NBI, NDVI, and Fv/Fm were largely uncorrelated with Ant and only weakly associated with Flv. Under subN, these physiological traits also showed weak negative correlations with plant biomass-related traits (Figure 2C). Yield-related traits were strongly and positively correlated with NUE-related traits, including CFr, NFr, E_N,y_, and NUE, as well as with Tb. They also showed weaker associations with other trait groups, being positively correlated with some physiological and root architecture-related traits and negatively correlated with C_N,y_ and Ant (Figure 2C).

### Multivariate trait responses to N availability

To identify the combination of traits most strongly associated with N availability while minimizing the influence of experimental design and spatial effects, Partial Least Squares Discriminant Analysis (PLS-DA) was performed using the adjusted phenotypic means estimated across the core subset of ToMAGIC lines. This analysis revealed a clear separation between plants grown under opN and subN conditions (Figure 3A). The first two latent components captured the major variation associated with N treatments, explaining 19% and 12.2% of the total variance, respectively (Figure 3A and 3B). When evaluated on an independent test set, the model achieved a classification accuracy, sensitivity, and AUC of 0.939, indicating strong discrimination between N treatments (Figure S2). The traits contributing most strongly to the separation along Component 1 were mainly agronomic and physiological traits. In particular, Ant traits were associated with the subN treatment, whereas shoot biomass-related traits were associated with the opN condition (Figure 3B). Component 2 mainly reflected variation in NUE- and yield-related traits (Figure 3B).

**Figure 3.**
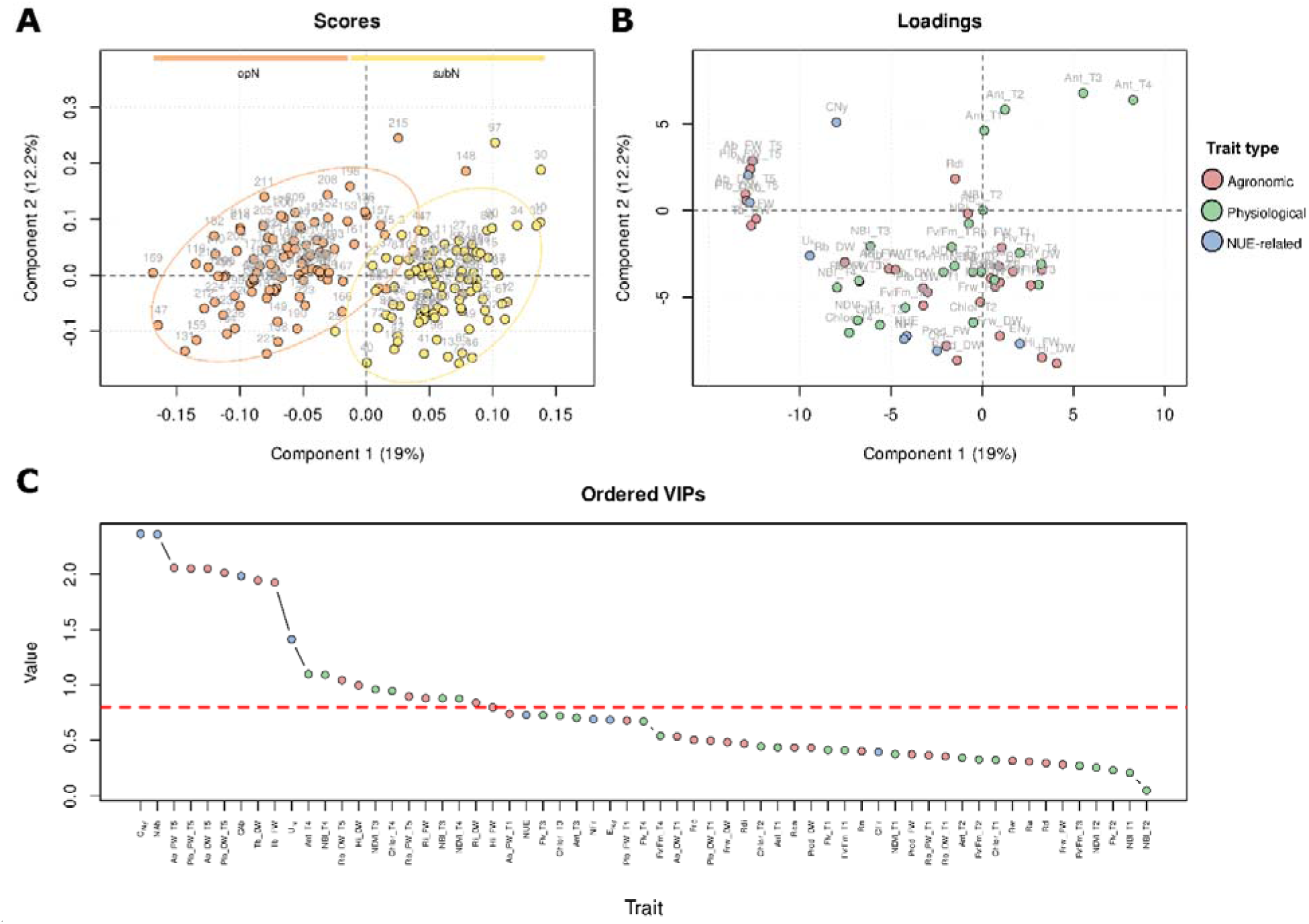
Partial least squares discriminant analysis (PLS-DA) of phenotypic traits measured in the core subset of ToMAGIC lines under optimal nitrogen (opN; 15 mM N) and suboptimal nitrogen (subN; 8 mM N) levels. **A.** Scores plot showing the separation of samples according to N treatment along the first two latent components. **B.** Loadings plot showing the contribution and direction of individual traits to treatment separation. Trait groups are indicated by color code, according to agronomic (red), physiological (green), and NUE-related (blue) traits. **C.** Ordered variable importance in projection (VIP) scores. The dashed red line indicates the threshold (VIP = 0.8) used to identify traits with strong discriminatory power.

Trait importance was further assessed using variable importance in projection (VIP) scores. Several traits showed substantial contributions to treatment discrimination (VIP > 0.8), including 11 agronomic traits measured at T5, six physiological traits measured at T3 and T4, and four NUE-related traits (Figure 3C). Most VIP-selected traits also corresponded to traits strongly affected by subN conditions, either positively or negatively (Figure 1B). However, some traits with smaller average responses, such as NDVI_T4, also showed VIP values above 0.8 (Figure 3C). This suggests that these traits provided complementary information for treatment discrimination, likely because they were less redundant with other selected traits (Figure 2C). Conversely, some traits that were markedly affected by N treatment did not reach the VIP threshold. For example, E_N,y_ increased by 42% under subN conditions (Figure 1B), yet its VIP value was 0.62. This lower VIP score likely reflects the strong correlation between E_N,y_ and other selected traits, such as Hi_DW (Figure 2C), which captured similar information in the model and reduced the relative contribution of E_N,y_ to class discrimination.

### Cross-environment consistency and reliability of genotype performance

Before defining proxy traits for NUE-oriented selection, we examined the consistency and reliability of predicted genotype performance across N treatments. This analysis was used to assess whether trait performance under opN could reliably inform selection under subN, or whether treatment-specific responses should be considered. Overall, empirical correlations between treatment-specific BLUPs varied markedly among traits, indicating different degrees of genotype ranking stability across N environments (Figure 4 and Figure S3). Yield-related traits, including Prod, Hi, Frw_FW, and Frc, together with NUE-related traits, including NFr, CFr, NUE, and E_N,y_, showed the highest consistency between treatments, with ρ ranging from 0.85 to 0.92 (Figure 4 and Figure S3). These traits also showed high prediction reliability (Table S9), supporting their use for selection across N conditions. By contrast, root-related traits, particularly root width (Rw), Rd, and Rdi, as well as NAb, showed lower correlations between treatments (ρ = 0.24–0.37), indicating stronger genotype-by-treatment changes and lower ranking stability (Figure S3). These traits also tended to show lower prediction reliability, suggesting that selection based on them should be interpreted with caution (Table S9).

**Figure 4.**
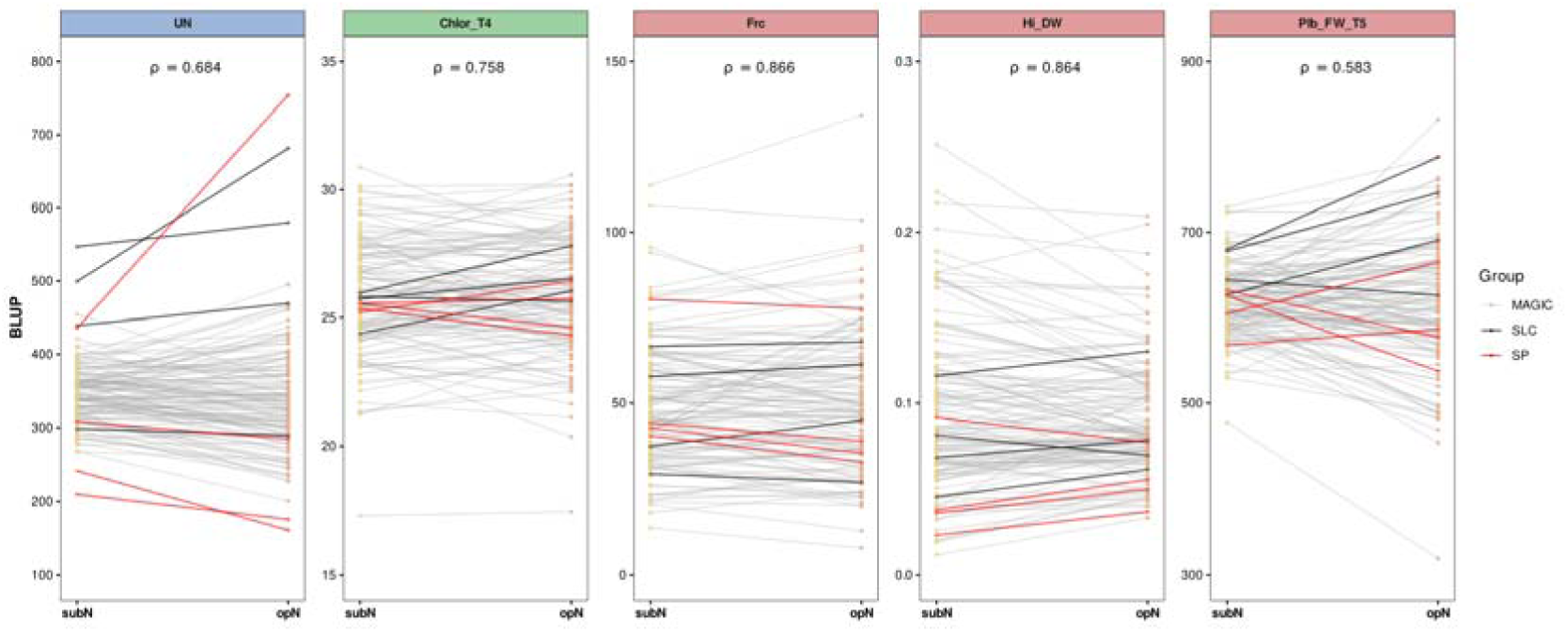
Reaction norms of predicted genotype performance across optimal nitrogen (opN; 15 mM N) and suboptimal nitrogen (subN; 8 mM N) levels for the proxy traits used in the multi-trait selection indices. Grey lines represent ToMAGIC lines, red lines represent SP founders, and black lines represent SLC founders. Best linear unbiased prediction (BLUP)-based correlations between treatments, estimated from the ToMAGIC lines, are shown in each panel.

### Selection of proxy traits for multi-trait NUE indices

Treatment response magnitude, heritability, and trait correlations were jointly considered to select a reduced set of informative traits for multi-trait selection. Traits with a VIP value above 0.80 were preferred (Figure 3C). In addition, traits with higher BLUP prediction reliability were prioritized, as they provide more robust genotype rankings for selection decisions (Table S9). Based on these criteria, U_N_, Chlor_T4, Frc, Hi_DW, and Plb_FW_T5 were selected as proxy traits representing complementary dimensions of N use, plant physiological status, fruit production, biomass partitioning, and plant vigor (Figure 4). U_N_ was selected as the main NUE-related proxy because it showed a strong negative response to subN (Figure 1B) and high heritability (h^2^ = 0.91 under opN and h^2^ = 0.77 under subN) (Table S6). In addition, U_N_ was slightly positively correlated with plant biomass traits (Figure 2C). Chlor_T4 was retained as a physiological indicator of plant status, as it was one of the most responsive physiological traits to subN level (Figure 1B) and also showed high heritability (h^2^ = 0.80 under opN and h^2^ = 0.83 under subN) (Table S6). Frc was included as a direct measure of fruit production. Although its VIP value was below the 0.8 threshold (VIP = 0.50), it showed a marked response to N treatment (Figure 1B) and high heritability (h^2^ = 0.87 under opN and h^2^ = 0.88 under subN) (Table S6). In addition, Frc showed strong correlations with NUE-related traits, including NUE, E_N,y_, NFr and CFr (Figure 2C). Finally, Hi_DW was selected as an indicator of biomass partitioning towards fruit production, whereas Plb_FW_T5 was included as a proxy for vegetative vigor. Hi_DW showed strong responses to N availability (Figure 1B) and high heritability (h^2^ = 0.88 under both opN and subN) (Table S6). Plb_FW_T5 was considered with reduced emphasis due to its moderate heritability under subN (h^2^ = 0.57), relatively high general prediction uncertainty (rSE = 0.65), and the need to avoid overvaluing excessive biomass accumulation relative to reproductive performance.

### Identification of high performance and subN-adapted lines

To identify promising breeding lines, the core subset of ToMAGIC lines was ranked using three multi-trait selection indices (SH, FAI_BLUP, and MGIDI) under two selection scenarios: high-performance lines and subN-adapted lines. All six index-by-scenario combinations produced positive total GG, ranging from 138% to 158% for the high-performance scenario, and from 125% to 170% for the subN-adapted scenario. In both cases, the SH index generated the lowest total GG, whereas the FAI-BLUP index generated the highest (Table 1). Hi_DW contributed most to the total GG, except for the subN-adapted SH index, where Frc showed the largest contribution. In contrast, Plb_FW_T5 exhibited negative GG after selection by MGIDI in both scenarios (Table 1).

**Table 1.**
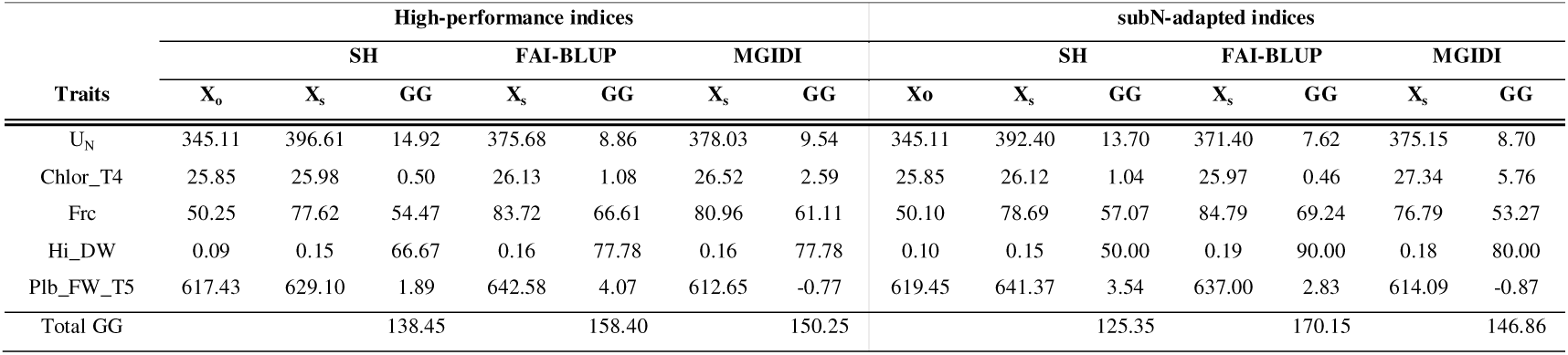
Genetic gains (GG) for the proxy traits used in the multi-trait indices under two selection scenarios: high-performance and subN-adapted lines. SH: Smith-Hazel index; FAI-BLUP: factor analytic best linear unbiased prediction index. MGIDI: multi-trait genotype-ideotype distance index. X_o_ and X_s_ represent the population mean before and after selection, respectively. GG indicates the percentage genetic gain for each trait.

The high-performance indices showed greater overlap among selected lines than the subN-adapted indices, resulting in 15 and 17 unique lines, respectively. Among these, lines 402, 428, 518, 800, and 816 were consistently selected across all six index-by-scenario combinations (Figure 5).

**Figure 5.**
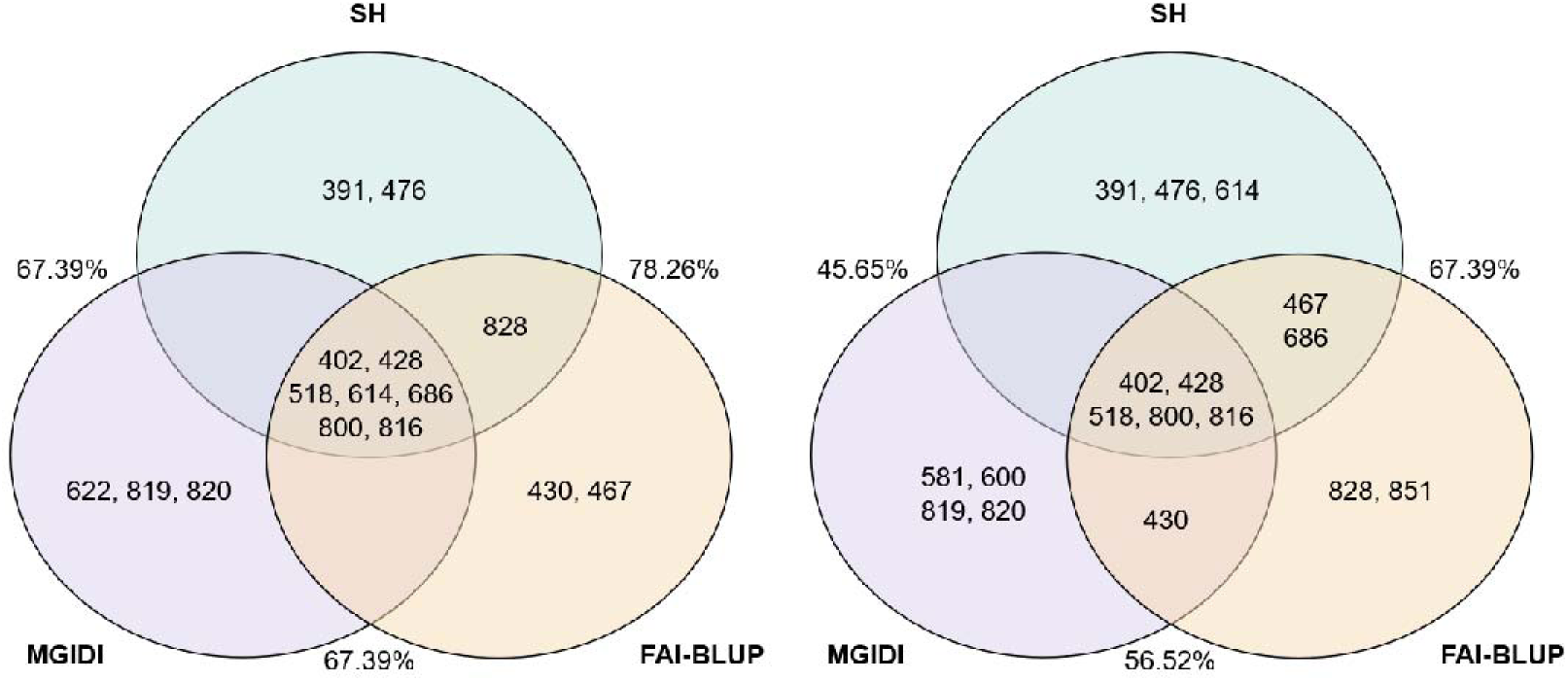
ToMAGIC lines selection by three multi-trait selection indices for high-performance (left) and subN-adapted (right) lines. SH: Smith-Hazel index; FAI-BLUP: factor analytic best linear unbiased prediction index. MGIDI: multi-trait genotype-ideotype distance index. Numbers inside the diagrams correspond to selected genotype identifiers. Percentages indicate the coincidence index (CI), representing the percentage of shared genotypes between pairs of indices.

The selected lines represented contrasting favorable responses. Line 428 exhibited the highest performance under opN, characterized by elevated shoot biomass and yield, as well as high NUE and related parameters. Although a substantial reduction in these traits was observed under subN conditions, this line still maintained comparatively high values relative to those of the collection (Table S3 and Table S10). Notably, a marked increase was detected in E_N,y_, reflecting enhanced N partitioning efficiency toward the fruit. Line 518 displayed lower biomass, yield, and NUE values than line 428 under opN. However, this line exhibited the highest tolerance to N limitation among the selected lines, showing the smallest reductions in key traits such as yield, NUE, E_N,y_, NFr, and CFr (Table S10). In contrast, lines 800 and 816 exhibited limited changes in biomass accumulation responses to reduced N availability (Table S10), predominantly at early developmental stages in the case of line 800, and during later stages associated with fruit development in line 816. Finally, line 402 showed vigorous biomass accumulation under opN (Ab_T5, Plb_T5, and NAb), together with a comparatively consistent response of root-related traits (Ri, Rd, and Rw) as well as U_N_ under subN (Table S10).

Root architectural variation among the selected lines was further assessed using the centroid displacement metric derived from the PCA space of the first three principal components, which accounted for 96.3% of the total variance (Figure S4). Lines 402, 428, 518, and 816 showed relatively small distances between opN and subN centroids, indicating limited multivariate shifts in root system architecture between N treatments. By contrast, line 800 showed a larger centroid displacement, ranking among the most responsive lines for root architecture changes under reduced N availability (Figure S4). Visual inspection of root images supported these patterns, with line 800 showing reduced rooting depth and a greater abundance of fine lateral roots under subN, while line 402 displayed visually thicker roots under opN than under subN (Figure S4). Because Rdi was not included in the PCA-based analysis, this difference was not reflected in the centroid displacement metric. Together, these observations suggest that maintenance of performance under reduced N supply can be achieved through both stable and highly responsive root phenotypes, indicating that multiple root-based strategies may contribute to NUE in the ToMAGIC population.

## Discussion

N fertilization has been a key driver of crop production over the past decades, enabling substantial increases in agricultural yields worldwide (Adalibieke et al., 2023). However, the environmental impacts associated with excessive N use have become a major concern, highlighting the need to reduce N inputs while maintaining crop productivity (Cao et al., 2023; Hu et al., 2023; Wu et al., 2023; Laidig et al., 2024). As observed in this study, reduced N availability affected cultivated tomato performance at multiple levels, including biomass accumulation, biomass partitioning, nutrient allocation, and fruit production (Sun et al., 2023). Therefore, identifying genotypes capable of maintaining good agronomic performance under limited N supply represents a major objective in modern breeding programs.

The broad phenotypic variability here observed across the ToMAGIC core subset is consistent with previous reports of NUE-related variation in tomato-related germplasm (Tripodi et al., 2022; Flores-Saavedra et al., 2024; Machado et al., 2024; Renau-Morata et al., 2024; Cirillo et al., 2025). However, from a breeding perspective, the value of this population lies in the incorporation of diversity from SLC and SP accessions into a structured and directly selectable genetic background (Arrones et al., 2024). In eggplant, studies using introgressions from wild relatives have shown that exotic diversity can generate wide phenotypic variation and transgressive individuals for yield, biomass, and NUE-related traits under low N conditions (Villanueva et al., 2021, 2026). This is relevant because the direct use of exotic or traditional materials can be limited by heterogeneous genetic backgrounds, limited agronomic performance, or linkage drag (Prohens et al., 2017). By reshuffling alleles from multiple founders across advanced fixed lines, MAGIC populations reduce the confounding effects of unstructured germplasm and enable the assessment of new allele combinations in a breeding-relevant context (Cavanagh et al., 2008; Huang et al., 2015; Arrones et al., 2020). Specifically, the ToMAGIC population represents a valuable resource, as it incorporates allelic diversity from SLC and SP founders while maintaining full sexual compatibility with cultivated tomato (Gramazio et al., 2020; Arrones et al., 2024). Its relevance for NUE-oriented breeding is further supported by the adaptive NUE responses previously reported in the founders under low N inputs (Flores-Saavedra et al., 2024; Renau-Morata et al., 2024; Gil-Villar et al., 2025). In the present study, the evaluated core subset of ToMAGIC lines showed broad phenotypic variation across agronomic, physiological, and NUE-related traits under opN and subN conditions. Importantly, the value of this population lies not only in the extent of its diversity but also in the agronomic relevance of the phenotypes observed, including transgressive lines for several traits. Thus, the evaluated ToMAGIC material provides a useful genetic resource for identifying novel allelic combinations and admixed genetic backgrounds associated with vigorous plant architecture, improved reproductive performance, and favorable NUE-related traits.

Beyond confirming the usefulness of the ToMAGIC core subset, our results provide insight into which traits may be most informative for NUE-oriented selection and breeding. This is important because NUE is a complex trait comprising three key components: N uptake, internal N utilization in biomass production, and N harvest index (Lammerts van Bueren and Struik, 2017). Accordingly, breeding for improved NUE requires the evaluation of both efficiency-related parameters and the agronomic and physiological traits that support them (Wang et al., 2022a; Mahboob et al., 2024; Natarajan et al., 2025; Zeng et al., 2026). Consistent with this multi-trait view of NUE, reducing N supply to 8mM triggered coordinated responses across biomass accumulation, production, resource partitioning, root development, and NUE-related traits. Notably, the reduction in aerial biomass was not accompanied by a proportional decline in fruit production, suggesting that the tested genotypes maintained reproductive performance by preferentially allocating available resources to fruits under subN conditions. This response contrasts with previous reports in cultivated tomato cultivars, including Moneymaker (Renau-Morata et al., 2024) and Ailsa Craig (Thompson et al., 2026), where reduced N fertilization typically decreased both vegetative biomass and fruit yield. Therefore, the partial uncoupling between biomass accumulation and fruit production in the ToMAGIC core subset suggests that genotypes differed not only in their sensitivity to subN treatment, but also in their capacity to acquire N and allocate it to reproductive growth. In this framework, U_N_ may represent a key integrative component linking above- and belowground responses to N limitation. Greater shoot vigor has been associated with enhanced N uptake under low N availability, as reported in wheat, where early vigor and shoot biomass strongly influenced N acquisition (Kamiji et al., 2014), while increased N uptake has also been linked with greater shoot biomass production in cowpea (Aliyu et al., 2023). At the same time, root architectural traits may support this uptake by improving soil exploration efficiency rather than simply increasing root biomass, consistent with the idea that specific root configurations enhance nutrient capture while minimizing the metabolic costs associated with root construction and maintenance (Lynch et al., 2024; Arrones et al., 2026).

Moreover, the largely conserved genetic correlation patterns among traits under opN and subN suggested that a substantial proportion of the genetic architecture underlying NUE-related traits remains stable across contrasting N conditions. This stability indicates that common genetic factors may coordinate trait relationships independently of N availability, a pattern also reported in other crop species, including rapeseed, rice, and barley (Bouchet et al., 2016; Vishnukiran et al., 2020; Zeng et al., 2024).

From a breeding perspective, phenotypic responsiveness alone is not sufficient to define reliable selection criteria. The extent to which trait variation is genetically controlled must also be considered. Indeed, indirect selection is expected to be more effective when target traits exhibit limited genetic variation, and the selected traits show a clear relationship with the target traits or breeding objectives (Alonso et al., 2018; Rajendran et al., 2022; Price et al., 2024). In our study, shoot biomass N content (NAb) was the trait most negatively affected by subN treatment, but it also showed one of the highest proportions of environmental variance, suggesting that its direct use as a selection criterion may be limited. Similarly, root system architecture traits showed a relatively low proportion of genetic variance and low-to-moderate heritability estimates compared with the other evaluated traits. Root traits have been associated with improved N uptake efficiency and overall NUE under N limitation in several crops (Ajmera et al., 2022; Jia et al., 2022; Wang et al., 2022b). However, their phenotyping remains challenging because root measurements are often destructive, labor-intensive, and highly sensitive to environmental and sampling variation. Therefore, the lower reliability observed for these traits in our study may partly reflect the difficulty of capturing root variation accurately under the present experimental conditions. Although biologically relevant, root traits may require more controlled or repeated phenotyping before being used as reliable direct selection targets in the ToMAGIC material. In contrast, several fruit- and NUE-related traits showed a predominance of genetic variance and consistently displayed high heritability, indicating strong genetic control and good potential for selection. Notably, many of these traits retained considerable genetic variance under subN conditions, suggesting exploitable variation for selection under N limitation.

The empirical genetic correlations between opN and subN estimated from the mixed model suggest a largely shared genetic basis across N treatments and limited genotype × N interaction. This interpretation is consistent with multi-environment breeding frameworks, in which high genetic correlations across environments indicate limited genotype re-ranking and support selection across environments (Bashir et al., 2023; Christensen et al., 2025; Lippolis et al., 2025). In this sense, selection under opN would be expected to identify genotypes with similar genetic potential under subN. This was reflected in the overlap between the two objective-driven selection indices, as some lines were selected for both high overall performance and specific adaptation to subN. However, this interpretation must be considered together with trait reliability. BLUPs may be affected by prediction error, shrinkage, low heritability, or high environmental variance, reducing reliability and making genotype rankings less stable (Togashi et al., 2018; Atanda et al., 2022; Yan et al., 2022). Therefore, traits with high biological relevance but low reliability may help describe the response to N limitation but should be used cautiously as primary selection criteria.

The use of multi-trait indices is consistent with breeding approaches that show selection is more effective when several traits are combined according to their genetic parameters, reliability, and relationships with the breeding objective (Li and Wu, 2023; Cole, 2025). In the present study, all index-by-scenario combinations resulted in positive genetic gains, supporting the suitability of the selected proxy traits for NUE-oriented selection. Across both breeding scenarios, Hi_DW and Frc contributed strongly to the predicted gains, reinforcing the importance of biomass partitioning toward reproductive organs under reduced N supply (Meng et al., 2021; Wang et al., 2022a). At the same time, the inclusion of U_N_ and Chlor_T4 further allowed the indices to account for N uptake and plant physiological status. The limited gain observed for Plb_FW_T5 reflected the lower emphasis deliberately assigned to vegetative vigor, preventing biomass accumulation from dominating the selection response. In addition, using different indices in parallel can strengthen genotype identification, as each index relies on different statistical assumptions and weighting strategies (Smith, 1936; Hazel, 1943; Rocha et al., 2018; Olivoto and Nardino, 2021). For instance, Ghazvini et al. (2024) identified two superior barley genotypes under drought stress using SH, FAI, and MGIDI indices, while Singh et al. (2024) selected four promising mustard genotypes for salt tolerance using the same approach. Similarly, Ambrósio et al. (2024) identified three early, high-yielding black bean genotypes that were consistently selected across four multi-trait indices. However, to our knowledge, this comparative framework has not previously been applied to NUE-oriented selection. In agreement with these studies, the overlap observed among the three indices used for each breeding scenario supports the robustness of the selected ToMAGIC lines. Lines 402, 428, 518, 614, 686, 800, and 816 were identified as high-performance lines, while 402, 428, 518, 800, and 816 were also selected as the most promising subN-adapted lines. This overlap suggests that these genotypes combine broad agronomic value with specific suitability under reduced N supply, making them promising candidates for NUE-oriented tomato breeding. Importantly, their favorable performance was not restricted to the proxy traits included in the indices. Most of these lines also showed values above, or close to, the population means for additional NUE-related traits, including NUE itself, its component traits, and C and N accumulation in shoot biomass and fruits. This broader response supports the biological consistency of the selected ideotypes and suggests that the indices captured relevant dimensions of N efficiency beyond the specific traits used for selection.

Interestingly, the complementary root analyses suggest that this favorable performance was not associated with a single root architectural response to N limitation. Instead, selected lines appeared to follow contrasting belowground strategies. Lines 428, 518, and 816 maintained relatively stable root architectures across N treatments, suggesting a capacity to sustain performance without major structural reorganization of the root system. By contrast, line 800 showed a stronger architectural adjustment under subN, indicating a more plastic response to reduced N availability. Together, these results indicate that improved adaptation to low N supply may be achieved through contrasting strategies, including either maintaining a stable root architecture or deploying a highly plastic root response, as also suggested by Sandhu et al. (2016) in rice and by Lecarpentier et al. (2021) in oilseed rape. The consistent identification of these lines across independent agronomic and morphometric analyses further supports their value as promising pre-breeding material for the development of NUE-adapted tomato cultivars.

## Conclusions

This study demonstrates the potential of the ToMAGIC population for NUE-oriented tomato breeding under contrasting N conditions. Broad phenotypic and genetic variation was detected and maintained under subN supply, including transgressive phenotypes for key productivity and NUE-related traits, providing a suitable basis for genomic selection. Although N limitation strongly affected plant performance, it mainly altered trait magnitude rather than the underlying genetic ranking of genotypes, supporting the feasibility of selection under both opN and subN environments. Importantly, unlike the responses commonly described in commercial tomato, the core subset of the ToMAGIC population largely maintained fruit production under subN conditions while increasing biomass and N partitioning towards fruits. This response highlights the potential of this germplasm to sustain reproductive performance under reduced N availability and represents a valuable target for breeding programs aimed at improving NUE without compromising productivity. The integration of agronomic, physiological, and NUE-related traits through complementary multi-trait selection indices enabled the identification of informative proxy traits and robust genotype selection for both high-performance and subN-adapted breeding objectives. In particular, ToMAGIC lines 402, 428, 518, 800, and 816 consistently combined favorable agronomic performance with improved adaptation to reduced N availability. These lines represent promising candidates for further validation and valuable pre-breeding material for developing tomato cultivars with improved performance under reduced N inputs. Together, these results support the use of structured multi-parent populations and multi-trait selection approaches to accelerate the development of tomato cultivars adapted to reduced N fertilization.

## Materials and methods

### Plant materials and experimental design

The ToMAGIC population, developed from the intercross of four *S. lycopersicum* var. *cerasiforme* (SLC) and four *S. pimpinellifolium* (SP) founders (Arrones et al., 2024), was used for this study. This population consists of 354 highly inbred lines, and a core subset of 118 lines was selected based on maximization of the genetic and phenotypic diversity using the Core Hunter 3 software (De Beukelaer et al., 2018). Genomic data from 6,488 high-quality SNP markers, along with six phenotypic traits including fruit, plant, and leaf morphology descriptors, were employed for line selection to capture the full diversity of the entire population (Arrones et al., 2024).

The eight founders and the core subset of 118 ToMAGIC lines were germinated in seedbeds with coconut fiber. Plantlets at the three-to-four-leaf stage (30-day-old) were transferred to 15 L pots filled with expanded clay balls (2–3 mm diameter; Arlita™, Madrid, Spain) and cultivated in a completely randomized design under greenhouse conditions at the Universitat Politècnica de València campus (39°28′55′′N, 0°20′11′′W). After acclimatization, 45-day-old plants were differentially fertigated and grown under two N supply conditions, ranging from opN (15 mM) to subN (8 mM), based on the Hoagland solution (Hoagland and Arnon, 1950). A total of 788 pots were arranged in 40 columns and 20 rows, with columns alternating between N treatments. Two plants were transplanted per pot, totalling 1,576 plants. Overall, six technical replicates were grown for each ToMAGIC line and treatment, randomly distributed across the experiment, while each founder was grown with 10 replicates per treatment. Half of the plants of each genotype and treatment were sampled after 15 days of differential fertilization (60 days-old plants; T1), retaining only one plant per pot, and the remaining half at the end of the experiment (135 days-old; T5). Plants were vertically trained with strings, and axillary buds were pruned weekly and kept free of insects and pests, following standard greenhouse management practices. Cultivation was continued to allow fruit development until the fourth truss.

### Genotypic data

Genotypic data for the founders and ToMAGIC lines, generated by Gramazio et al. (2020) and Arrones et al. (2024), respectively, were used in this study. To expand the latter subset of 6,488 SNPs, candidate SNPs for the founders and ToMAGIC lines underwent separate filtering rounds. For founders, the final set of high-quality SNPs was obtained by selecting biallelic SNPs with distinct genotypes between the eight accessions using BCFtools and requiring support from at least 10 reads (--minDP 10) using VCFtools v. 0.1.16 (https://vcftools.github.io/man_latest.html). For the ToMAGIC lines, candidate SNPs were initially filtered by missing data (--max-missing 0.8), genotype and site mean depth (--minDP 3 and --min-meanDP 10), and alternative allele frequency (--maf 0.05) using VCFtools. Additionally, a LD k-nearest neighbour genotype imputation method (LD KNNi) was applied to fill the missing calls or genotyping gaps (Troyanskaya et al., 2001). Only common SNPs between the founders and ToMAGIC lines sets were retained, resulting in 10,684 SNP markers.

### Phenotypic data

A total of 29 primary traits were measured throughout the experiment, including 15 agronomic traits, 6 physiological traits, and eight NUE-related parameters (Table S1). Plants were cut and partitioned into Ab and Rb, and measured at two time points: after 15 days of differential fertilization (T1) and at the end of the experiment (T5). These traits were assessed at both FW and DW. Additionally, roots were cleaned and photographed, and different root system architecture traits were calculated using GiA Roots software (Figure S5; Galkovskyi et al., 2012). Fruits were harvested at the red-ripe stage throughout the experiment. At T5, the cumulative harvest was used to determine Prod, Frc, and Frw. Physiological traits were repeatedly measured at four time points (T1–T4) in one leaf per plant (the fifth or sixth leaf from the apex), with measurements spaced 15 days apart. Repeated measurements of the same trait were treated as separate variables in the analysis. The total shoot biomass and fruit content of elemental C and N was determined by the Dumas combustion method using a TruSpec Micro CHNS elemental analyser (LECO Corp, Michigan, USA) at the Servicio de Análisis de C y N of the Estación Experimental del Zaidín (CSIC, Granada, Spain). The NUE and its components, including U_N_ as the ratio between mean plant N content during the mean growth period and N content in the seed, E_N,y_ as the ratio between fruit production and the mean plant N content during the mean growth period, and C_N,y_; were calculated following the methodology of Weih (2014), which was optimized for tomato by Domínguez-Figueroa et al. (2020), being NUE = U_N_ x E_N,y_ x C_N,y_.

### Statistical analysis

Treatment-specific genetic parameters were estimated using the sommer v4.4.4 package (Covarrubias-Pazaran, 2016) to fit a diagonal multi-treatment mixed model of the form:

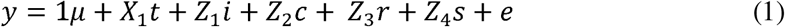

where y is the vector of phenotypic data; µ is the overall mean; t is the vector of fixed effect of the nitrogen treatment; i is the vector of random genotype effects structured by treatment, where i ∼ N(0, S⊗G); c is the vector of random effect of the column, where c ∼ 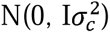; r is the vector of random effect of the row, where r ∼ 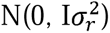; s is the vector of a 2D-spline random effect to capture finer-scale spatial trends not explained by row and column effects, where s ∼ 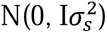; and e is the vector of random residuals, where e ∼ N(0, R). G is the additive genomic relationship matrix constructed by the Gmatrix function from the AGHmatrix v2.1.4 R package with the VanRaden parameter (Amadeu et al., 2023). S and R are structured diagonal matrices that allow estimation of different genetic and residual variances for each treatment (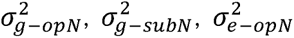 and 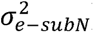). *X*_1_, *Z*_1_, *Z*_2_, *Z*_3_, and *Z*_4_ are the incidence matrices of their respective effects.

The residuals were examined for normality using diagnostic plots after model fitting. Outliers were identified as residuals exceeding three times the standard deviation of the standardized residual. The significance of the fixed effects was assessed using the Wald test, while the significance of the random effects was evaluated using the likelihood-ratio test between the full model in Equation (1) and a reduced model without the effect to be tested.

After outlier removal, the model was re-fitted to estimate general and treatment-specific best linear unbiased predictions (BLUPs) and the narrow-sense heritability (h^2^) for each treatment using the following formulas:

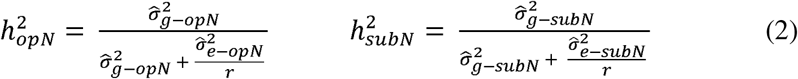

where r is the number of replications.

Prediction uncertainty was summarized using the standard error (SE) of the BLUPs. For general BLUPs, relative SE was calculated as the mean prediction SE divided by the standard deviation of the predicted values. For treatment-specific BLUPs, prediction error variance (PEV) was calculated as SE^2^, and prediction reliability was estimated as:

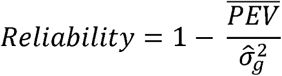

where 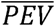 is the mean prediction error variance and 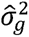 is the treatment-specific additive genetic variance estimated from Equation (1).

Descriptive statistics were then calculated from the cleaned phenotypic dataset for each trait, separately for the ToMAGIC core subset, SLC founders, and SP founders under opN and subN conditions. These included range, mean, median, and CV. Differences among genetic groups within each N treatment were tested using a linear mixed model fitted with the lme4 v1.1-37 R package (Bates et al., 2015), with genetic group included as a fixed effect and genotype as a random effect. Differences between N treatments within each genetic group were assessed using a second linear mixed model including the genetic group × N treatment interaction as a fixed effect and genotype as a random effect. Post hoc pairwise comparisons were performed using estimated marginal means with the emmeans v2.0.0 R package, applying Tukey adjustment for multiple comparisons (Lenth, 2025). Significance was declared at p < 0.05. Pairwise Pearson’s correlation coefficients among traits were then estimated separately within each N treatment.

To estimate adjusted means for each genotype under each treatment, a model similar to Equation (1) was fitted, with genotype and genotype-by-treatment interaction treated as fixed effects. Exploratory empirical genotypic correlations among traits and between treatments were calculated as Pearson correlations using treatment-specific BLUPs.

### Partial least squares discriminant analysis (PLS-DA)

PLS-DA was performed to identify the traits that best discriminated between N levels using genotype-level adjusted means. N treatment was used as the response variable, and BLUEs of the phenotypic traits were used as predictors. Missing values were imputed using a random forest approach implemented in the rfImpute function of the randomForest v4.7-1.2 R package (Liaw and Wiener, 2002), and variables were centered and scaled prior to analysis. The model was fitted in R using the mdatools v0.14.2 package (Kucheryavskiy, 2020), with 10-replicated 5-fold cross-validation. The number of components was selected according to the Wold’s criterion. Trait importance was assessed using VIP scores, considering variables with VIP > 0.8 as relevant (Wold et al., 2001). Model predictive performance was evaluated through misclassification rate, sensitivity, specificity, and the area under the curve (AUC) from corresponding receiver operating characteristic (ROC) curves.

### Selection of promising lines with multi-trait indices

The selection indices were used to rank genotypes according to two breeding objectives: (I) the identification of high-performance lines with good overall agronomic performance, with particular emphasis on performance under subN, and (II) the identification of lines specifically adapted to subN, defined as genotypes with good production under subN, independently of their performance under opN. Three multi-trait selection indices were evaluated (Supplementary Methods): the Smith-Hazel (SH) index, the factor analytic best linear unbiased prediction (FAI-BLUP) index, and the multi-trait genotype-ideotype distance index (MGIDI), all implemented in the metan R package v1.19.0 (Olivoto and Lúcio, 2020). Because these indices differ in their mathematical formulations and in how they account for trait relationships and ideotype definition, they were applied in parallel to provide complementary perspectives for genotype ranking.

To construct selection indices that were both biologically meaningful and operationally useful for breeding, it was necessary to identify a reduced set of traits representative of NUE and plant performance. Trait selection was therefore guided by five criteria: (i) magnitude of the response to N availability, (ii) heritability estimates, (iii) correlation structure among traits to avoid redundancy, (iv) contribution to treatment discrimination in the PLS-DA analysis (VIP scores), and (v) genotype performance reliability. The vector of BLUPs for the selected traits, obtained from the model in Equation 1, varied depending on the selection objective. For the high-performance selection index, subN BLUPs were used for U_N,_ and Chlor_T4 to capture stress-related variation in NUE and N status, while general BLUPs were used for Frc, Hi_DW, and Plb_FW_T5 to represent overall fruit production, biomass partitioning, and plant vigor. For the subN-adapted index, subN BLUPs were used for all selected traits in order to prioritize genotypes with superior performance under low-input agriculture.

### Estimation of the coincidence index (CI) and the genetic gains (GG)

To evaluate the performance of the different multi-trait indices, the top 10 ranked lines were selected from the total set of 118 lines, corresponding to a selection intensity of 8.5%. This selection intensity was used to compare the indices’ ability to identify superior genotypes under the two breeding objectives and to estimate the coincidence index (CI) and the genetic gains (GG).

Selection efficiency was examined via the CI between each pair of selection indices as described by Hamblin and Zimmermann (1986):

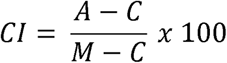

where *A* is the number of selected genotypes common to different methods, *C* is the number of expected genotypes selected by chance, and *M* is the number of genotypes selected according to the selection intensity.

The predicted GG, which expressed the genetic superiority of the selected lines, was primarily assessed using the BLUPs values as:

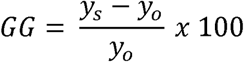

where *y_s_* was defined as the mean breeding value of the selected genotypes and *y_o_* was defined as the mean breeding value of the population.

### Root system architecture analysis

To summarize multivariate variation in RSA, four representative traits (Ra, Rd, Rw, and Rn) were subjected to principal component analysis (PCA). The first three principal components were retained and used to represent every individual in a reduced multivariate space. For each genotype and N treatment, centroid coordinates were calculated as the mean PC1–PC3 scores of all individuals belonging to that genotype-treatment combination. Euclidean distances between opN and subN centroids were subsequently computed and used as an integrated measure of RSA responsiveness to N availability, with larger distances indicating greater treatment-induced architectural changes. Centroid estimation and distance calculations were performed using a custom Python script developed for this study and available from the authors upon request.

## Acknowledgments

This work has been funded by MICIU/AEI/10.13039/501100011033, by the European Union NextGenerationEU/PRTR (TED2021-129296B-I00), MICIU/AEI/10.13039/501100011033 (PID2020-118627RB-I00), MICIU/AEI/10.13039/501100011033 and ERDF/EU (PID2022-13654IOB-I00), Conselleria d’Innovació, Universitats, Ciència i Societat Digital from Generalitat Valenciana (CIPROM/2021/020 and CIAICO/2023/167), and European Union Horizon 2020 research and innovation programme under grant agreement No 101000716 (HARNESSTOM project). V.B.-F. is grateful to Conselleria d’Educació, Cultura, Universitats i Ocupació (Generalitat Valenciana), for a predoctoral contract (CIACIF/2023/238). AA is grateful to Conselleria d’Educació, Cultura, Universitats i Ocupació (Generalitat Valenciana) for a postdoctoral grant (CIAPOS/2024/330). Pietro Gramazio is grateful to MICIU/AEI/10.13039/501100011033 and the European Union through NextGenerationEU/PRTR for a post-doctoral grant (RYC2021-031999-I). Funding for open access charge: Universitat Politècnica de València.

## Author contributions

Conceptualization: RVM, SGN, JP. Methodology: VB-F, JB, MSJ, MP, PG, AG, RVM, AA. Investigation: VB-F, DG-V, JB, BR-M, MSJ, AG, AA. Resources: SV, JMP-P, AG, SGN, JP. Data curation: VB-F, DG-V, MSJ, SGN, AA. Formal analysis: VB-F, DG-V, JB, BR-M, PG, SV, JMP-P, JP, AA. Visualization: VB-F, DG-V, MSJ, MP, PG, SV, RVM, AA. Writing—original draft: VB-F, AA. Writing—review & editing: all authors. Supervision: SGN, JP, AA. Project administration: SGN, JP. Funding acquisition: JMP-P, AG, SGN, JP, AA.

## Supplementary information

**Table S1. List and description of agronomic, physiological, and nitrogen use efficiency (NUE)-related traits evaluated in the study.**

**Table S2. Raw phenotypic data for the 118 evaluated ToMAGIC lines and the eight founder accessions, including four S. lycopersicum var. cerasiforme (SLC) and four S. pimpinellifolium (SP), under optimal (opN; 15 mM) and suboptimal (subN; 8mM) N conditions.** Each row corresponds to an individual experimental unit and includes experimental design information, genotype identity, nitrogen treatment, replicate, and trait measurements.

**Table S3. Descriptive statistics of the evaluated traits in the core subset of ToMAGIC lines, *Solanum lycopersicum* var. *cerasiforme* (SLC), and *S. pimpinellifolium* (SP) founder accessions under optimal nitrogen (opN; 15 mM) and suboptimal nitrogen (subN; 8 mM) conditions.** For each trait and N treatment, minimum, mean ± standard deviation (SD), median, maximum, and coefficient of variation (CV, %) are shown. Different letters indicate significant differences among groups within each trait and nitrogen treatment based on pairwise comparisons of estimated marginal means from a linear mixed model using Tukey adjustment (p < 0.05). Asterisks indicate significant differences between opN and subN within each group and trait based on Tukey-adjusted pairwise comparisons from a separate linear mixed model (* p < 0.05, ** p < 0.01, *** p < 0.001).

**Table S4. Transgressive segregation detected in the ToMAGIC core subset.** The Summary sheet reports the frequency and magnitude of positive and negative transgression for each phenotypic variable, while the Lines sheet identifies the transgressive genotypes and their corresponding transgression values.

**Table S5. Significance of the fixed effects according to Wald test statistics and significance of the random effects according to the likelihood-ratio test.** *, **, *** significant at the 0.05, 0.01, and 0.001 probability levels, respectively.

**Table S6. Variance components and narrow-sense heritability (h^2^) estimates for the evaluated traits under optimal nitrogen (opN; 15 mM) and suboptimal nitrogen (subN; 8 mM) conditions.** Variance components were estimated using the multi-treatment mixed model and include treatment-specific additive genetic variance (σ^2^_gt-opN_and σ^2^_gt-subN_), row variance (σ^2^_r_), column variance (σ^2^_c_), 2D-spline variance (σ^2^_s_), and treatment-specific residual variance (σ^2^_e-opN_ and σ2_e-subN_).

**Table S7. Phenotypic trait correlations under optimal nitrogen (opN; 15 mM) and suboptimal nitrogen (subN; 8 mM) conditions.** Correlations under subN are shown in the lower triangle, whereas correlations under opN are shown in the upper triangle. Traits are ordered according to hierarchical clustering from the correlation opN heatmap.

**Table S8. Empirical genotypic trait correlations based on best linear unbiased predictions (BLUPs) under optimal nitrogen (opN; 15 mM) and suboptimal nitrogen (subN; 8 mM) conditions.** Correlations under subN are shown in the lower triangle, whereas correlations under opN are shown in the upper triangle. Traits are ordered according to hierarchical clustering from the correlation opN heatmap.

**Table S9. Prediction uncertainty and reliability of BLUPs for the evaluated traits under optimal (opN; 15 mM) and suboptimal (subN; 8 mM) N conditions.** Mean BLUPs and their standard errors (SE) were estimated for each trait from Equation (1). For general performance, uncertainty was expressed as relative SE (rSE). For treatment-specific BLUPs, prediction error variance (PEV) and reliability were estimated.

**Table S10. Descriptive statistics of the consistently selected high-performance ToMAGIC lines across all six index-by-scenario under optimal nitrogen (opN; 15 mM) and suboptimal nitrogen (subN; 8 mM) conditions.** For each trait and N treatment, mean ± standard deviation are shown.

**Figure S1. Phenotypic trait correlations under optimal nitrogen (opN; 15 mM) and suboptimal nitrogen (subN; 8 mM) supply.**

**Figure S2. Performance assessment of the partial least squares discriminant analysis (PLS-DA) model used to discriminate optimal nitrogen (opN; 15 mM) and suboptimal nitrogen (subN; 8 mM) treatments.** Misclassification rate, sensitivity, and specificity are shown across increasing numbers of latent components for calibration (cal) and cross-validation (cv). The receiver operating characteristic (ROC) curve summarizes model performance, with an area under the curve (AUC) of 0.939, indicating strong discrimination between both opN and subN conditions.

**Figure S3. Reaction norms of predicted genotype performance across optimal nitrogen (opN; 15 mM) and suboptimal nitrogen (subN; 8 mM) levels.** Grey lines represent ToMAGIC lines, red lines represent SP founders, and black lines represent SLC founders. Best linear unbiased prediction (BLUP)-based correlations between treatments, estimated from the ToMAGIC lines, are shown in each panel.

**Figure S4. Three-dimensional sample space of five selected lines under optimal nitrogen (opN; 15 mM) and suboptimal nitrogen (subN; 8 mM) treatments.** Five genotypes are shown in the principal component space (PC1–PC3), where each triangle represents three biological replicates per line and condition. Centroids (red dots) summarize the mean position of each condition, and arrows indicate the displacement between subN and opN, reflecting the magnitude of the RSA phenotypic response to treatment. Four of the five lines (#402, #428, #518, and #816) show minimal displacement between conditions, indicating low sensitivity to the treatment, whereas line #800 exhibits a pronounced shift, consistent with a strong response. Representative root images under subN and opN are also included, illustrating the comparison between both conditions.

**Figure S5. Representation of the root system architecture traits computed using Gia Roots software (Galkovskyi et al., 2012).** Root area (Ra), depth (Rd), and width (Rw) were measured from the pixels in the image that correspond to the entire root system. Root convex area (Rca) is defined as the area of the convex hull that contains the image corresponding to the entire root system. Average root diameter (Rdi) was measured as the mean value of the root width estimation computed for all pixels of the medial axis of the entire root system. The maximum number of roots (Rn) was measured as the 84^th^ percentile value after sorting the number of roots crossing a horizontal line from lowest to highest.

