## Supplementary Methods for "Multi-trait evaluation of a tomato MAGIC population identifies promising lines with improved nitrogen use efficiency (NUE)"

*Smith-Hazel (SH) index*

The SH index (Smith, 1936; Hazel, 1943) integrates the phenotypic and additive genetic variance-covariance matrices to maximize the expected genetic response, using predefined economic weights assigned to each trait. The selection index values were calculated as:

$b=P^{-1}Ga$	(3)

where $P$ and $G$ denote the phenotypic and additive genetic covariance matrices, respectively, and $a$ is the vector of economic weights. Since the direct economic values of the traits were difficult to quantify, economic weights were defined using an iterative procedure (Jahufer and Casler, 2015). Multiple combinations were evaluated, and the resulting selection responses were examined. The final weighting scheme was chosen as the one producing a pattern of predicted genetic gains consistent with the breeding objectives: 3 for U_N_ and Chlor_T4, 1 for Frc and Hi_DW, and 0.1 to Plb_FW_T5.

*Factor analytic best linear unbiased prediction (FAI-BLUP) index*

The FAI-BLUP index was estimated following Rocha et al. (2018) using factor analysis and an ideotype design. Unlike covariance-based linear indices, the FAI-BLUP approach accounts for the correlation structure among traits and explicitly specifies the desired direction of selection for each trait. Here, we selected the maximum value as the desirable ideotype for each trait, except for Plb_FW_T5, for which the mean value was considered. Factor analysis was applied to the predicted genetic values of the selected traits to obtain factor scores, after which the genotype-ideotype distance was calculated using standardized Euclidean distance and converted into a similarity probability:

$$P_{ij}= \frac{\frac{1}{d_{ij}}}{\sum_{i=1;j=1}^{i=n;j=m} \frac{1}{d_{ij}}}$$

where $P_{ij}$ was the probability of the $i^{th}$ ($i=1, 2, \ldots, n)$ genotype to be similar to the the $j^{th}$ ($j=1, 2, \ldots, m)$ ideotype; $d_{ij}$ was the genotype-ideotype distance between the $i^{th}$ genotype and the $j^{th}$ ideotype, calculated based on the standardized mean Euclidean distance.

*Multi-trait genotype-ideotype distance index (MGIDI)*

The MGIDI is based on the distance between each genotype and the ideotype and was estimated as described by Olivoto and Nardino (2020). This index first rescales all traits according to the desired direction of selection, so that favorable values are expressed on a common scale. We defined the maximum value as the desirable ideotype for each trait, except for Plb_FW_T5, for which the mean value was considered. Factor analysis was then applied to the rescaled data to account for trait correlations and reduce dimensionality. The MGIDI value for each genotype was subsequently calculated as the Euclidean distance between the genotype and the ideotype in the reduced factor space:

$${MGIDI}_{i}= \sqrt{\sum_{j=1}^{f} {(F_{ij}-F_{j})}^{2}}$$

where ${MGIDI}_{i}$ was the distance between the $i$th genotype and the ideotype, $F_{ij}$ was the score of the $i$th genotype in the $j$th factor, $F_{j}$ is the ideotype score for the $j$th factor, and $f$ is the number of retained factors.
